# *Toxoplasma gondii* transcription factor AP2XII-8: a key regulator of G1 phase progression and parasite division

**DOI:** 10.1101/2024.09.13.612814

**Authors:** Yuehong Shi, Qiang Yang, Yue He, Xingju Song, Dandan Hu

## Abstract

The apicomplexan parasite *Toxoplasma gondii* can infect humans and virtually all warm-blooded animals worldwide, posing a significant threat to public health and being of veterinary importance. Acute infections are characterized by the fast replication of tachyzoites inside host cells. During this fast amplification process, gene expression is highly regulated by a series of regulatory networks. The G1 phase, which is usually conserved across species, is responsible for preparing the materials necessary for the next replicating cell cycle; however, few regulators have been identified at this stage. Here, we functionally characterized the C/G1 phase-expressed ApiAP2 transcription factor, TgAP2XII-8, in *T. gondii* tachyzoites. Conditional knockdown of TgAP2XII-8 leads to significant growth defects and asexual division disorders. Additionally, parasite cell cycle progression was disrupted following TgAP2XII-8 depletion, characterized by G1 phase arrest. RNA-seq and CUT&Tag experiments revealed that TgAP2XII-8 acts as an activator of ribosomal proteins expressed in the G1 phase. Moreover, TgAP2XII-8 binds to a specific DNA motif ([T/C]GCATGCA), which is abundant and conserved in the intergenic region of several other apicomplexans, possibly suggesting a broad and conserved role for this ApiAP2 in the Phylum of Apicomplexa. Our study reveals that TgAP2XII-8 acts as a critical C/G1 phase regulator, orchestrating the cell cycle in *T. gondii* tachyzoites. This study contributes to a broader understanding of the complexity of the parasite’s cell cycle and its key regulators.

## 1 Introduction

*Toxoplasma gondii* is an obligate intracellular parasitic protozoan known for its intricate life cycle that encompasses multiple hosts and developmental stages. The life cycle of *T. gondii* can be categorized into sexual and asexual reproduction. The sexual phase is restricted to the intestinal epicellular cells of definitive hosts within the Felidae family, while the asexual phase occurs in any warm-blooded intermediate host, including humans[1]. In immunocompetent hosts, acute infections are usually asymptomatic, whereas in immunocompromised individuals, such as those with AIDS, fetuses carried by pregnant women, and those with compromised immune systems, they can lead to severe diseases and even fatalities[2]. During the asexual phase, rapid generation of numerous tachyzoites is facilitated by endogenous action. In response to host immune pressure and other factors, tachyzoites can differentiate into bradyzoites as a strategy to withstand external stresses. Upon the relief of these pressures, the bradyzoites have the ability to reactivate into tachyzoites, thereby causing acute toxoplasmosis[3].

*T. gondii* has the ability to sense alterations in its external environment. It adeptly modulates its gene expression status in a timely manner, thereby regulating the progression of the cell cycle to accommodate changes in its growth metabolism. During the asexual reproductive phase, the cell cycle of *T. gondii* can be categorized into G1 (Gap 1), S (DNA synthesis), G2 (Gap 2, which was previously presumed to be missing), M (Mitosis), and C (Cytokinesis or budding) phases [4, 5]. *T. gondii* undergoes replication within the intermediate host (tachyzoites, bradyzoites) through a process known as endodyogeny[6, 7]. During endodyogeny, the cell nucleus initially undergoes duplication and then splits into two equal parts under the influence of the centrosome. At the same time, the action of the inner membrane complex (IMC) segregates the two cell nuclei and organelles, gradually forming two progeny cells while dismantling the mother cell. Notably, the cell cycles of tachyzoites within the same parasitophorous vacuole are typically synchronized, revealing a precise and robust regulatory system to coordinate the synchronized division of these parasites[4]. Transcriptomic analysis of synchronous cell cycle populations revealed that approximately 40% of genes exhibit a cyclic pattern. This suggests that *T. gondii* may rely on a ’just-in-time’ expression pattern to generate transcripts and proteins as needed[8]. Moreover, in classical eukaryotic cells, the cell cycle is intricately regulated by a complex network involving cell checkpoints, intricate protein phosphorylation and protein degradation[9, 10]. It is believed that a restriction checkpoint exists during the G1 phase, which receives external signals and determines whether the organism proceeds to the division cycle or not[11]. In contrast to tachyzoites, *T. gondii* bradyzoites feature an extended G1 phase, creating a differentiation window that provides sufficient time for the shift in transcriptional expression profiles and metabolic changes between tachyzoites and bradyzoites[12]. While the actual commitment to the transition from tachyzoites to bradyzoites may occur during the S/M phase, the decision is initiated during the G1 phase[13].

The timing of gene expression during the cell division cycle is intricately regulated by transcription factors[14–16]. Apicomplexan parasites have a unique family of transcription factors characterized by the possession of one or more plant-like AP2 DNA-binding domains. These factors are known as ApiAP2 transcription factors, which bind to specific promoter sequences and regulate the expression of target genes[17]. A total of 68 ApiAP2s have been identified in *T. gondii*, of which 24 are considered to be cyclically expressed ApiAP2s[8]. Functional studies have revealed that ApiAP2s are involved in the regulation of gene expression at different developmental stages and cell cycles of parasites. TgAP2XI-4, TgAP2IX-4, TgAP21b-1, TgAP2IV-3, TgAP2IV-4, and TgAP2IX-9 have been established as regulators of bradyzoite formation[18–22]. Recent studies have further demonstrated the involvement of TgAP2XII-2, TgAP2XII-1 and TgAP2XI-2 in gene regulation and developmental transitions during merozoite, pre-sexual and sexual stages[23–25]. TgAP2XI-5 exhibits consistent expression throughout the tachyzoite cell cycle and participates in the regulation of virulence genes and other gene expression during the S/M phase by binding to the GCTAGC motif and synergizing with TgAP2X-5[26].

In this study, we functionally characterized the C/G1 phase-expressed ApiAP2 transcription factor TgAP2XII-8. Our findings reveal its role in targeting multiple untranslated regions of *T. gondii* genes and activating nuclear ribosomal protein expression during the G1 phase. Knocking down TgAP2XII-8 results in significant growth defects and disruption of parasite cell cycle progression, characterized by G1 phase arrest, which potentially affects the coordination of parasite replication, as its absence leads to asynchronous replication and endodyogeny. These data provide basic knowledge for understanding the transcriptional regulation of parasite cell cycle in *T. gondii*.

## 2 Materials and methods

### 2.1 Parasites and cell culture

*T. gondii* RH ΔKu80 strain and its derivative strains were continuously cultured in vitro with human foreskin fibroblasts (HFFs; ATCC, Manassas, VA, USA) or African green monkey kidney (Vero) cells using Dulbecco’s Modified Eagle’s Medium (DMEM, Macgene, Beijing, China) supplemented with 10% heat-inactivated fetal bovine serum (TransGen Biotech, Beijing, China). The cells were cultured at 37°C in a 5% CO_2_ atmosphere.

#### Generation of transgenic *T. gondii* strains

The guide RNA targeting TgAP2XII-8 was designed using the EuPaGDT library from the ToxoDB database. The CRISPR-Cas9 plasmid and donor fragments were constructed following the strategy presented in Figure 1D. For the Cas9-expressing plasmid, the upstream and downstream fragments featuring Cas9 and gRNA sequences were amplified and joined using the Seamless Cloning Kit (Vazyme Biotech, Co, Ltd, Nanjing). To construct the TgAP2XII-8-mAID plasmid, the mAID sequence and 3×HA epitope tag were fused to the C-terminus of TgAP2XII-8. The plasmid (mAID-3HA-CAT-TIR1-3Flag) was amplified and ligated to the chloramphenicol resistance gene (CmR) and the TIR1-3Flag expression cassette. For the C-terminal epitope tagging of TgAP2XII-8, a 59bp PCR primer was developed from the mAID-3HA-CAT-TIR1-3Flag plasmid for PCR amplification of the donor, containing a 39bp fragmented upstream of the TgAP2XII-8 translational termination codon and downstream of the gRNA site. The resulting homologous recombinant PCR fragment was used to co-transfect the RH ΔKu80 strain and its corresponding CRISPR-Cas9 plasmid, and screened for parasites in a medium containing 20 μm chloramphenicol. Monoclonal parasites were identified using PCR and IFA techniques. At a particular culture stage, the degradation of TgAP2XII-8 was induced by IAA at a final concentration of 500 μM. All primers utilized in the experiment can be found in Supplementary table 1.

**Figure 1.**
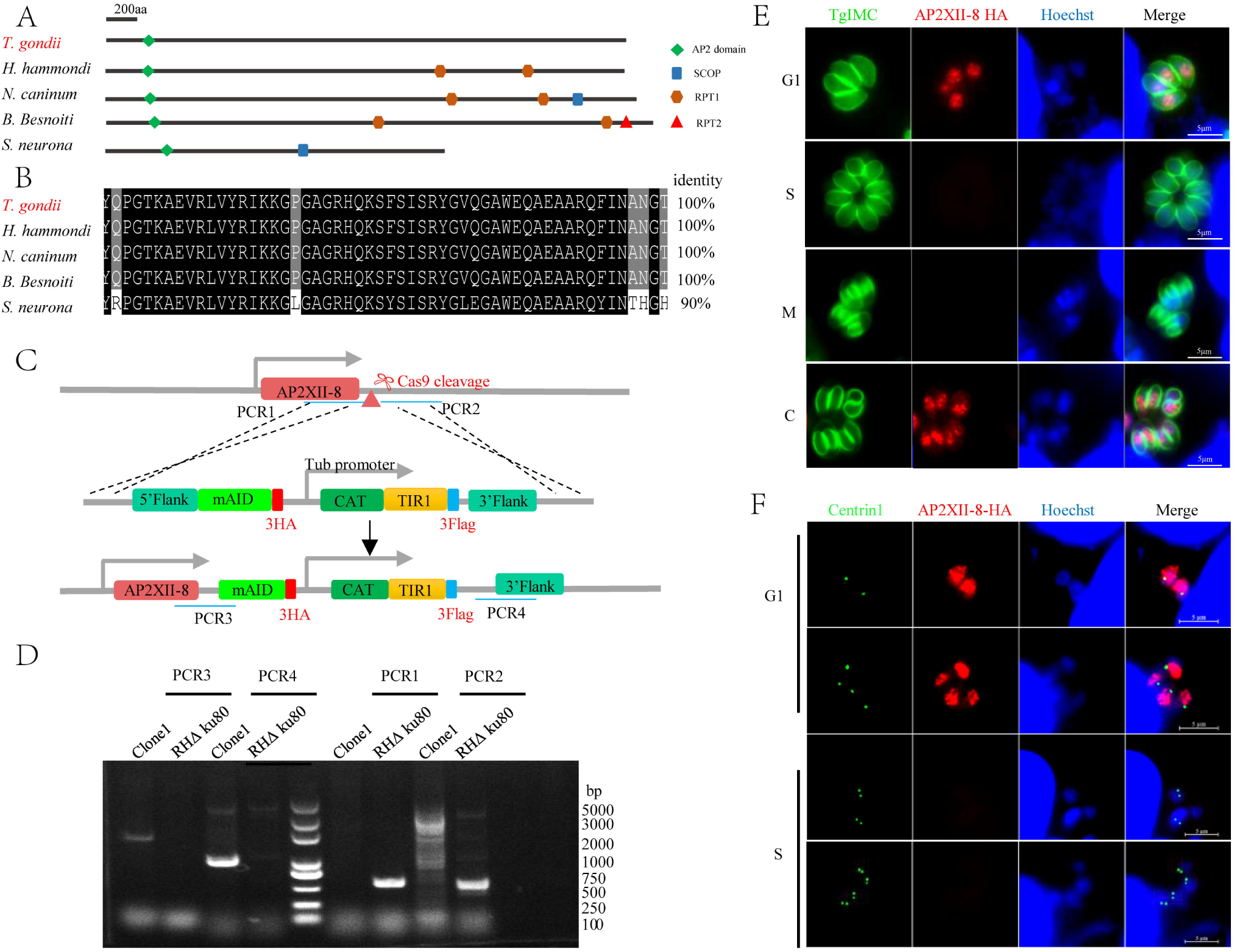
TgAP2XII-8 is expressed in the C and G1 phases of *T. gondii.* (A) Schematic representation of the conserved domain of TgAP2XII-8 (TGGT1_250800) and its orthologs from other organisms, including *Hammondia hammondi* (HHA_250800), *Neospora caninum* (NCLIV_066340), *Besnoitia besnoiti* (BESB_029280) and *Sarcocystis neurona* (SN3_01600620). Domains were predicted by SMART[68]. (B) Multiple sequence alignment of AP2 domains. The AP2 domain sequences from TgAP2XII-8 were aligned with their homologs. Regions of high identity and similarity between AP2 domain sequences are shown as black and gray columns. The percentage of homology between TgAP2XII-8 and each AP2 domain is shown at the end of the alignment. (C) Diagram showing the strategy for tagging TgAP2XII-8 with AID-3xHA in wild parasites. The location of primers used to integrate the PCR is indicated. (D) PCR demonstrating homologous integration and gene tagging in representative clones compared to the RH strain. (E-F) TgAP2XII-8 protein expression during the tachyzoite cell cycle. Co-immunostaining of intracellular tachyzoites was performed using mouse anti-HA (red), rabbit anti-inner membrane complex 1 (TgIMC1, green), or rabbit anti-TgCentrin 1 (green) antibodies. Hoechst 33258 was used for nuclear staining. TgIMC1 staining was employed to detect budding daughter parasites. Since TgIMC1 staining patterns do not precisely distinguish between G1 and S phases, these phases are identified by TgCentrin1 staining (F). Parasites in the G1 phase containing a single centrosome, whereas those in the S phase exhibit duplicated centrosomes. Cell cycle stages are shown on the left side of the images: G1, gap phase; S, synthesis phase; M, mitotic phase; C, cytokinesis.

### 2.2 Immunofluorescence assay (IFA)

Intracellular immunofluorescence experiments were performed according to previous studies[27]. Briefly, freshly harvested parasites were inoculated onto HFF cells grown on glass coverslips in 12-well plates. After incubation, infected cells were properly fixed with 4% PFA for 30 min, permeabilized with 0.25% Triton X-100, and then blocked with 3% bovine serum albumin (BSA) for 30 min. The samples were incubated with mouse anti-HA (1:500, Sigma, St. Louis, MO, USA), mouse anti-TY (1:500), rabbit anti-TgGAP45 polyclonal antibodies (1:300), rabbit anti-TgIMC1 polyclonal antibody (1:300), and rabbit anti-TgCentrin 1 antibody (1:500) for 1 h. The samples were washed three times with PBS and then incubated with FITC-or Cy3-conjugated secondary antibodies (1:100, Proteinnovogtech, Rosemont, IL, USA) for 1 h. Nuclear DNA was stained with Hoechst 33258 (1:100, Macgene, Beijing, China). Polyclonal antibodies to TgGAP45 and TgIMC1 were validated in our previous study[28] and were gifts from Professor Qun Liu, China Agricultural University; mouse anti-TY and rabbit anti-TgCentrin 1 antibodies were gifts from Professor Shaojun Long, China Agricultural University[29]. Images were obtained using a Zeiss Fluorescence Microscopy system (Zeiss, Germany).

### 2.3 Intracellular replication assay

The intracellular replication ability of *T. gondii* tachyzoites was evaluated by IFA as previously described[27]. HFF cells in 12-well plates were infected with 1 × 10^5^ fresh RH tachyzoites per well and incubated for 1 h. The cell surface was then washed twice with PBS to remove extracellular tachyzoites, followed by incubation with IAA (500 μM) or vector (1:1000 ethanol). After 24 h of treatment, cells were fixed with 4% PFA and then immunofluorescence staining was performed using rabbit anti-TgGAP45 polyclonal antibodies (1:300) and Hoechst33258 (1:100, Macgene, Beijing, China). The number of parasites per strain was determined by counting at least 100 vacuoles using fluorescence microscopy. Three biological replicates were assessed to determine the number of tachyzoites per PV.

### 2.4 Invasion assay

Parasites, grown for 48 h with or without IAA, were harvested. Purified fresh tachyzoites, suspended in DMEM medium with or without IAA, were inoculated onto HFF monolayers and incubated at 37°C for 1 h. The coverslips were then fixed with 4% PFA followed by immunofluorescence staining. Extracellular parasites were stained with mouse anti-TgSAG1 polyclonal antibody (1:300) and mouse FITC-conjugated secondary antibodies (1:100, Proteintech, Rosemont, IL, USA). The cells were then permeabilized with 0.25% TritonX-100, and all parasites were stained with rabbit anti-TgGAP45 antibody and rabbit Cy3-conjugated secondary antibodies (1:100, Proteintech, Rosemont, IL, USA). The invasion efficiency was detected by two-color staining. Three biological replicates were assessed, and at least 15 random fields were counted for each replicate.

### 2.5 Egress Assay

Parasites were inoculated into 12-well plates and cultured for 30 h with or without IAA treatment. The egress was triggered with 3 μM of Ca^2+^ ionophore A23187 (Macklin, Shanghai) for min at 37□ before fixation with 4% PFA[27, 30]. IFA was performed using rabbit anti-TgGAP45 antibodies. A total of 100 vacuoles were randomly selected to count the ruptured vacuoles/whole vacuoles per slide. Three independent experiments were performed.

### 2.6 Plaque assay

Briefly, HFF cells growing in 12-well plates were infected with 100 freshly harvested tachyzoites and incubated for 7 days without disturbance with vector (EtOH 1:1000) or IAA (500 μM) containing medium. Then, the infected HFFs were fixed with 4% paraformaldehyde and observed by staining with 0.2% crystal violet solution. The plaque area was counted by pixel using Photoshop C6S software (Adobe, United States), and data were compiled from three independent experiments.

### 2.7 RNA-Seq and data analysis

The iKD TgAP2XII-8 strain cultured in Vero cells was treated with either 500 μM IAA or vehicle for 12 and 24 h, respectively. Total RNAs from freshly harvested tachyzoites were then extracted using the M5 Total RNA Extraction Reagent (Mei5 Biotechnology Co., Ltd, Beijing). Each treatment consisted of three biological replicates. The purity, concentration and integrity of RNAs were tested using the NanoPhotometer® (IMPLEN, CA, USA), the Qubit® RNA Assay Kit in Qubit® 2.0 Fluorometer (Life Technologies, CA, USA), and the RNA Nano 6000 Assay Kit of the Bioanalyzer 2100 system (Agilent Technologies, CA, USA), respectively. Illumina sequencing libraries were generated using the NEBNext® Ultra™ RNA Library Prep Kit for Illumina® (NEB, USA) according to the manufacturer’s recommendations. Sequencing was performed using the Illumina Novaseq 6000 platform (Beijing Novogene Technology Co., Ltd) to generate 150 bp paired-end reads. The original sequencing data can be found in the Sequence Read Archive database under the accession number PRJNA1148922.

Data analysis was performed via an online pipeline on the BMKCloud (www.biocloud.net) platform based on the reference genome of *Toxoplasma* Type II ME49 strain (ToxoDB-57). Briefly, Paired-end clean reads were aligned to the reference genome using Hisat2[31]. Read counts were calculated for each gene using the sorted bam files from StringTie[32]. Differentially expressed genes (DEGs) between treated and untreated parasites were calculated by edgeR[33]. Gene expression with a fold change >2 or < -2 and FDR < 0.05 was considered to be significantly differentially expressed. TPM (Transcripts per kilobase million) values were calculated for each gene and used to generate clustered heatmaps.

### 2.8 CUT&Tag and data analysis

Fresh tachyzoites of HA-tagged TgAP2XII-8-mAID strain or RH strain (1 x 10^7^) were harvested after 24 h of incubation in 75 cm^2^ cell culture flasks. Library construction was performed using the NovoNGS CUT&Tag 4.0 High-Sensitivity Kit for Illumina (Novoprotein, Suzhou, China) according to the manufacturer’s instructions. Briefly, fresh tachyzoites were bound to activated concanavalin A beads (10 μL/sample) and incubated for 10 min at room temperature.

The mixture was resuspended and incubated with primary antibody (1:50, mouse anti-HA) at 4°C overnight. After several washes, the parasites were incubated with secondary antibody (1:100, goat anti-mouse IgG) for 1□h at room temperature. The parasites were then resuspended with pAG-Transposome buffer and incubated for 1 h at room temperature on a rotator. Tagmentation was stopped by MgCl_2_ treatment and DNA extraction was performed using DNA extraction beads (Novoprotein, Suzhou, China). According to the manufacturer’s recommendations, Illumina sequencing libraries were generated by PCR amplification using specific adaptors (NovoNGS CUT&Tag 4.0 High-Sensitivity Kit for Illumina B box, Novoprotein, Suzhou, China). The CUT&Tag libraries were sequenced using the Illumina Novaseq 6000 platform (Beijing Novogene Technology Co., Ltd).

The paired-end reads were filtered and then aligned to the *T. gondii* ME49 reference genome using Bowtie2[34] (v.2.1.0). The unmapped reads and PCR duplicates were removed from the sorted bam files using samtools[35] and Picard tools (https://broadinstitute.github.io/picard/), respectively. The filtered reads were then employed to identify CUT&Tag peaks using MACS2[36]. The overlapped peaks in the two biological replicates were identified by the Irreproducibility Discovery Rate (IDR) [37]. Final peaks were annotated to *T. gondii* protein coding genes according to the distance of each peak to the closest TSS using ChIPseeker[38]. The sorted and filtered bam files of CUT&Tag peaks and RNA-seq reads were normalized to RPKM with a resolution of 10 bp (bin size) and transformed into bigwig files for direct visualization in IGV (Integrative Genomics Viewer)[39]. The DNA binding motifs were analyzed with peaks at DNA fragments ≤ 600 bp to increase the sequence proportion containing the motif by using both MEME[40] and HOMER (http://homer.ucsd.edu/homer/) algorithms. DNA motif occurrences on the whole genome of *T. gondii* were identified using FIMO online software (https://meme-suite.org/meme/tools/fimo). Raw sequencing data and processed data are available in the NCBI GEO database under the accession number GSE275112.

### 2.9 Statistical analysis

Violin charts, line drawings, scatter plots, and histograms were generated using GraphPad Prism 9 (San Diego, CA, USA). Heatmaps were drawn using the OmicStudio tool[41] on https://www.omicstudio.cn/tool. All experiments were performed in independent biological replicates as described for each experiment in the manuscript. Statistical significance in plaque assays, invasion, proliferation, and parasite growth inhibition assays was evaluated by two-tailed unpaired *t*-tests or two-way ANOVA using GraphPad Prism. Statistical data are expressed as mean value ± standard error.

## 3 Results

### 3.1 TgAP2XII-8 is mainly expressed during the C/G1 phase of the cell cycle of tachyzoites

Several transcriptional regulators were previously predicted based on their cell cycle-regulated expression patterns[8]. Among these regulators, the focus was on TgAP2XII-8 (TGME49_250800), a transcript that is dynamically expressed with peak expression during the C/G1 phase[8](Figure 1A, toxodb.org). A sequence analysis of TgAP2XII-8 was first performed, with our results showing that TgAP2XII-8 contains one conserved AP2 domain located at the 115-158 amino acids of its N-terminus. The AP2 domain of TgAP2XII-8 shares over 80% amino acid homology with other apicomplexans, including *Hammondia hammondi* (*HHA_250800*), *Neospora caninum* (NCLIV_066340), *Besnoitia besnoiti* (BESB_029280) and *Sarcocystis neurona* (SN3_01600620) (Figure 1A and B). To confirm that the TgAP2XII-8 protein is expressed during the cell cycle, a mAID-3xHA epitope tag was fused to the C-terminal end of TgAP2XII-8 in the RH parasite strain by a CRISPR-Cas9 strategy (Figure 1C)[42]. The correct integration locus in the engineered strain was confirmed by PCR (Figure 1D). Co-staining using cell cycle markers, including *T. gondii* Inner membrane complex-1 (TgIMC1) [43] and Centrin-1 (TgCentrin-1) [44], demonstrated that TgAP2XII-8 is a nuclear protein mainly expressed in Cytokinesis (C) and G1 phases, as shown by co-localization of the HA staining signal with that of Hoechst staining (Figure 1E). Furthermore, TgAP2XII-8 was detected only in the G1 phase with a single centrosome, but not in the S phase with a double centrosome, by co-staining using HA and TgCentrin-1 antibodies (Figure 1F). This suggests that TgAP2XII-8 is no longer expressed or detected when the parasite progresses to the S/M phase (as indicated by early budding).

### 3.2 Conditional knockdown of TgAP2XII-8 blocks parasite replication and causes G1 phase arrest

To estimate the role of TgAP2XII-8 during the parasite lysis cycle, the indole-3-acetic acid (IAA) degradation (mAID) conditional gene knockdown system was employed (Figure 1C). This study demonstrated that TgAP2XII-8 may be depleted after treatment with IAA (Figure 2A). A plaque assay measuring the comprehensive ability of the parasite to grow over 7 days confirmed the absence of a lysis plaque phenotype in the TgAP2XII-8-mAID strain treated with IAA, in contrast to the distinct plaques observed in the untreated control (Figure 2B and C). These data suggest that TgAPXII-8 is essential for the lytic cycle and parasite growth.

**Figure 2.**
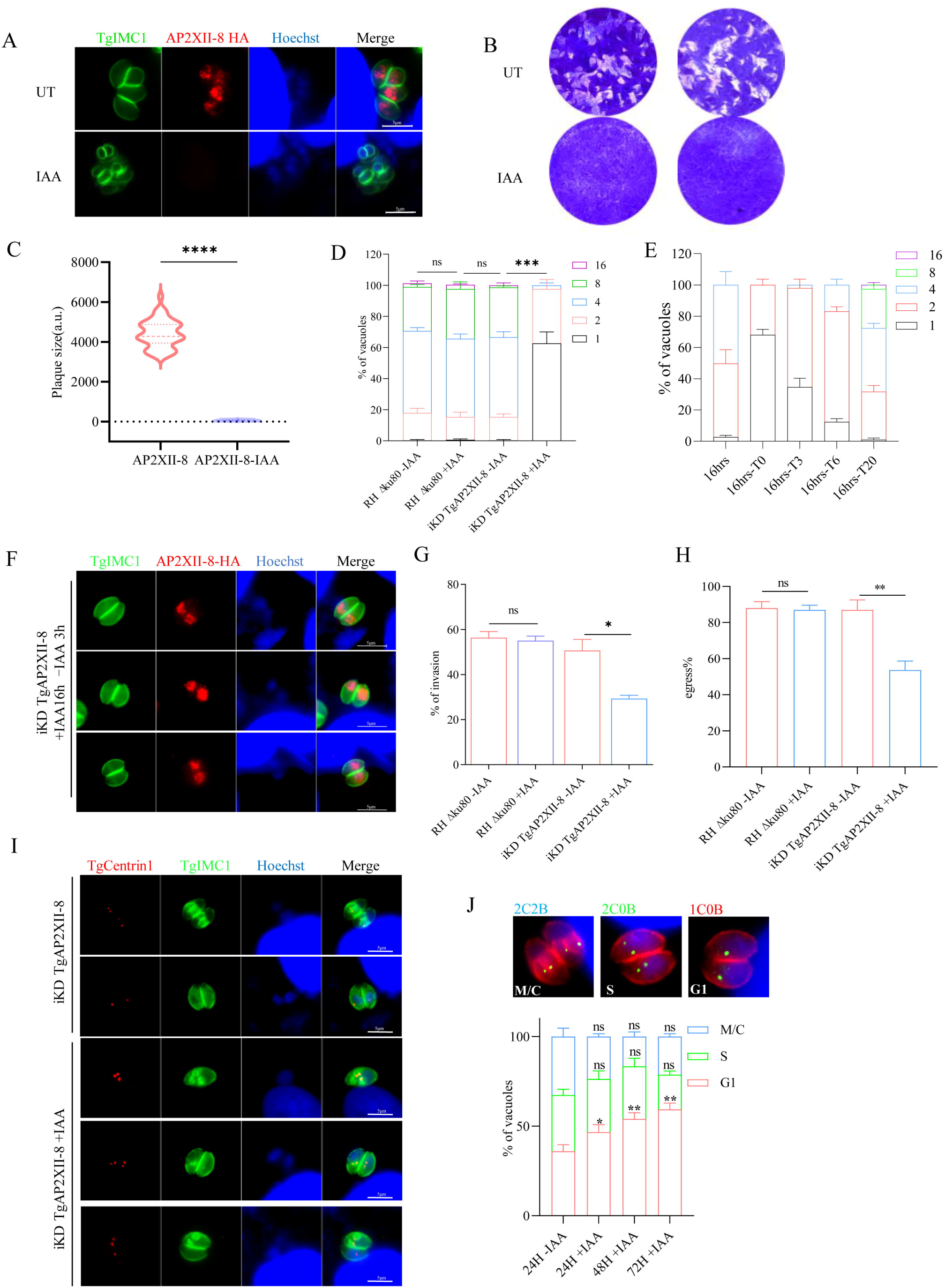
TgAP2XII-8 is essential for the proliferation of *T. gondii* tachyzoites. (A) Immunofluorescence assay of intracellular TgAP2XII-8-mAID parasites after treatment with IAA (500 µM) or vehicle (EtOH, 1:1000). Scale bars = 5 µm. (B) Plaque assay for the TgAP2XII-8 strain in the presence of IAA treatment or vehicle treatment for 7 days. (C) Measurements of plaque sizes in Photoshop C6S software (Adobe, USA) and expressed in pixel units. Over 30 plaques were analyzed for each repeat. (D) Intracellular replication assay. Parasites were incubated with IAA or vector at 24 h post-infection, and parasite replication was observed by IFA using anti-TgGAP45 antibody staining. Data are expressed as mean ± SEM of three independent assays, each with 100 total PVs calculated per strain. (E, F) Assessment of intracellular growth of iKD TgAP2XII-8 parasites after 16 h of culture in IAA-added medium, followed by culturing for 0, 3, 6, and 20 h in standard medium without IAA (16h-T0/T3/T6/T20). Parasite nuclei were stained with hoechst33258 and the parasite inner membrane complex (IMC) was stained with TgIMC1 antibody. The number of tachyzoites in the PV was counted by three independent assays of 100 PV each. (G) Invasion efficiency of TgAP2XII-8-mAID tachyzoites pretreated with or without IAA for 48 h. HFF cells were then infected at 37°C for 1 h. Data represent the mean ± SD of three independent experiments, each counting 15 fields per strain. (H) Efficiencies of A23187-induced egress of iKD TgAP2XII-8 parasites pretreated with or without IAA for 30 h. The average number of ruptured PVs was determined by randomly counting 100 vacuoles per slide. Mean ± SD of three independent experiments are graphed. (I) Observation of centrosome division by IFA in iKD TgAP2XII-8 parasites treated with or without IAA. Red, anti-TgIMC1; green, anti-TgCentrin1; Hoechst was used to stain the nucleus. Scale bars = 5 µm. (J) Quantification of the parasite cell cycles after IAA treatment for different times. Parasites in the G1 phase contain single centrosomes without budding (1C0B), whereas parasites in the S-phase have duplicate centrosomes without budding (2C0B), and parasites in the M/C phase have duplicate centrosomes with budding (2C2B). Centrosomes were detected using anti-TgCentrin 1 (green), and IMC was stained as red. At least 100 vacuoles were counted per biological replicate. Results are shown as mean ± SD of three independent experiments. *: *P* < 0.05, **: *P* < 0.01, ***: *P* < 0.001, ****: *P* < 0.0001, n.s: no significant difference.

Reduction in plaque formation may be caused by impairment of one or more steps of the lytic cycle, including invasion, intracellular replication, and egress. To determine which step of the parasite lytic cycle is affected by TgAP2XII-8 deletion, we conducted assays using the AP2XII-8 mAID strain to investigate these phenotypes. The intracellular replication ability of the parasite was observed in the presence of IAA or vehicle (ethanol [EtOH]), and TgAP2XII-8 was found to cause severe defects in replication in vitro (Figure 2D). TgAP2XII-8-mAID parasites treated with IAA for 24 h contained only 1-2 parasites per vacuoles. However, more vacuoles containing 4 and 8 tachyzoites were observed in the control. After removal of IAA from the culture, the parasite’s intracellular replication was significantly recovered over time, suggesting that deletion of the TgAP2XII-8 protein leads to reversible defects in parasite replication (Figure 2E and F). In addition, the invasion ability was also found to be significantly reduced after the conditional knockdown of TgAP2XII-8 (29%, *p*<0.05) (Figure 2G). For the egress assay, A23187-stimulated iKD TgAP2XII-8 parasites from HFF monolayers showed a significant decrease in the egress ratio (53.5%, *P*<0.001) after 30 h of treatment with or without IAA (Figure 2H). These results suggest that TgAP2XII-8 plays a key role in the lytic cycle of tachyzoites.

To further dissect the cause of slow replication in TgAP2XII-8 deletion mutants, we further probed replication defects by centrosome counting. Immunofluorescence assays (IFAs) with specific antibodies for centrosomes (Anti-TgCentrin 1) and subcellular budding (Anti-TgIMC1) were used to detect the percentage of parasites at different cell cycle stages[45]. Parasites with a single centrosome and no daughter buds were in the G1 phase, parasites with replicating centrosomes and no daughter buds were in the S phase, and parasites with replicating centrosomes and budding daughter were in the cytoplasmic division (C) phase. Among the parasites expressing TgAP2XII-8, approximately 36% were in the G1 phase, 31% in the S phase, and 33% in the C phase, whereas 47% of the accumulated parasites were still in the G1 phase after 24 h of IAA treatment. The G1-phase delayed parasites showed more enrichment after extended period of IAA exposure, reaching 54% and 59% after 48 h and 72 h of IAA treatment (Figure 2I and J). These results support a delay in the G1 phase of the cell cycle after TgAP2XII-8 deletion.

### 3.3 TgAP2XII-8 depletion results in subcellular organelle gene alterations

Given the potential transcription factor properties of TgAP2XII-8, an RNA-seq was used to examine changes in gene expression following depletion of the TgAP2XII-8 protein. TgAP2XII-8-mAID tachyzoites were treated with or without IAA for 12 h and 24 h respectively, and differentially expressed genes (DEGs) were identified with a minimum fold change of 2 and an adjusted *P* < 0.05. The results showed that a total of 357 genes were downregulated and 662 genes were upregulated (Figure 3A) at 12 h of IAA treatment, whereas a total of 748 genes were downregulated and 709 genes were upregulated (Figure 3B) at 24 h of treatment.

**Figure 3.**
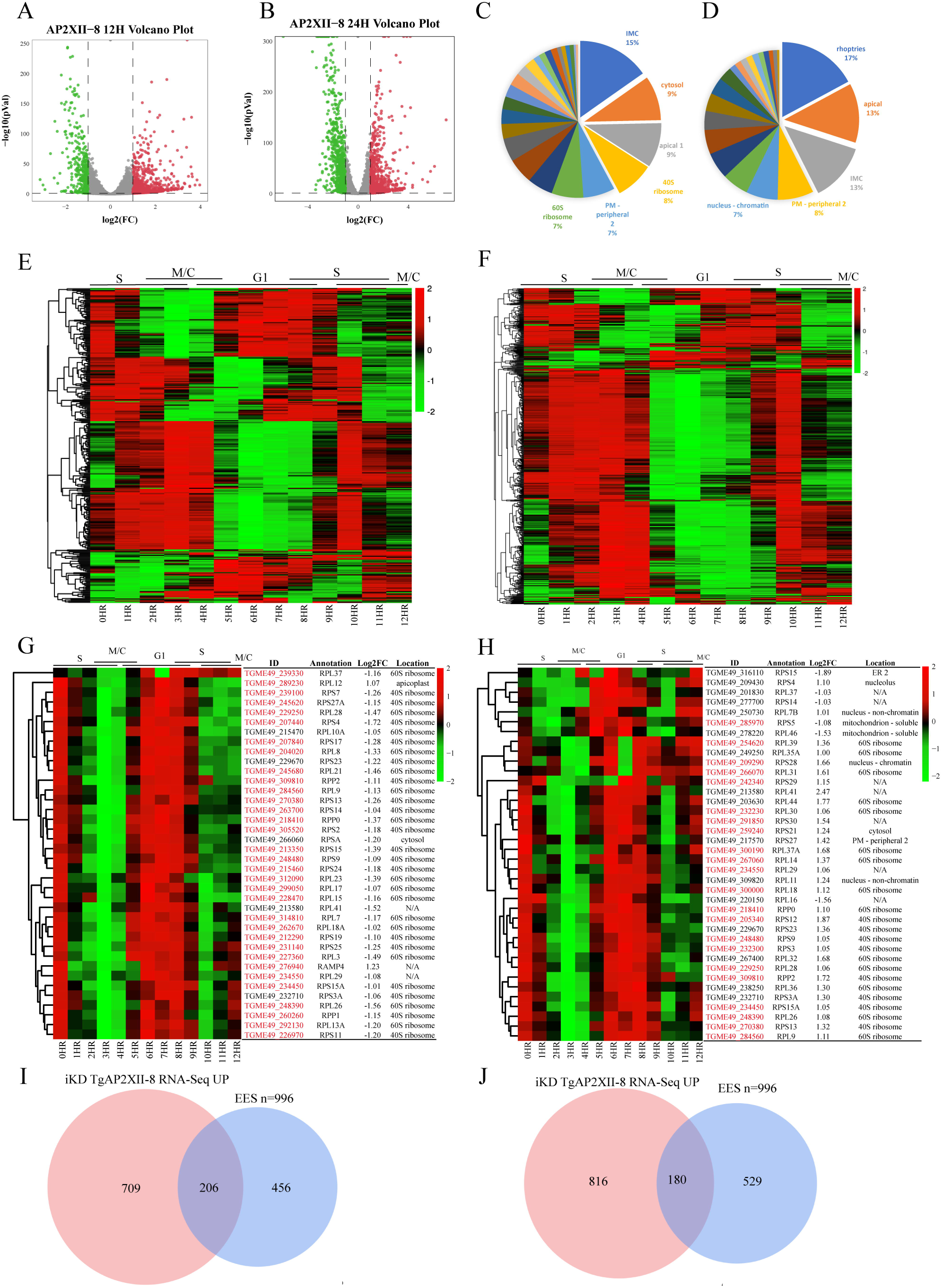
TgAP2XII-8 regulates the transcription of multiple cell-cycle genes. (A-B) Volcano plot of differentially expressed genes in TgAP2XII-8 parasites treated with IAA for 12 h (A) or 24 h (B) compared to untreated control by RNA-seq. Downregulated genes are shown in green and upregulated genes are shown in red. Statistically nonsignificant genes are indicated in gray. Differential expression analysis was based on three independent biological experiments. Predicted localization of downregulated genes after 12 h (C) and 24 h (D) of IAA treatment was assigned according to the hyperLOPIT proteomic dataset[46]. (E, F) Heatmaps showing the cell cycle expression of downregulated DEGs in the iKD TgAP2XII-8 strain after being treated with IAA for 12 h (E) and 24 h (F). The cell cycle gene expression data are referenced to Behnke et al.[8], with different phases (G1-S-M-C) indicated at the top. (G, H) Heatmaps showing cell cycle expression of ribosome proteins in the iKD TgAP2XII-8 strain after being treated with IAA for 12 h (G) and 24 h (H). CUT&Tag hits are marked in red. (I, J) Venn diagrams for the upregulated genes in 12 h (I) and 24 h (J) IAA-treated parasites and enteroepithelial stage-specific genes. For enteroepithelial stage-specific genes, see Ramakrishnan et al. (2019)[69].

To comprehensively assess the potential localization and function of the transcripts, the DEGs were mapped to the hyperLOPIT proteomic dataset[46]. Among the 357 downregulated genes in the TgAP2XII-8-mAID parasite after 12 h of IAA exposure, 32 ribosomal proteins from 40S ribosome and 60S ribosome as well as 58 IMCs and apical complex proteins were identified based on the hyperLOPIT proteomic dataset[46] (Figure 3C, Supplementary table 2). After 24 h of IAA exposure, more than 300 organelle proteins were downregulated, including rhoptry (75), IMCs (54), apical complex (55), apicoplast (19), mitochondria (31), ER & Golgi (37), and dense granule (20) (Figure 3D, Supplementary table 3). Almost all of these organelle genes showed downregulation, except for 46 dense granule proteins that were up-regulated. Notably, 22 ribosomal proteins were also found to be upregulated after 24 h of IAA treatment, even they were downregulated after 12 h of treatment (Supplementary table 3).

Since TgAP2XII-8 is cyclically expressed during the lytic cycle, the cell cycle expression patterns of all DEGs after TgAP2XII-8 depletion were examined using the transcriptome data of Behnk et al. [8]. The expression of the downregulated genes after 12 h and 24 h of IAA treatment was noted to occur mainly in the S and M-C phases, whereas they showed low expression in the G1 phase (Figure 3E and F), suggesting a high degree of consistency in the expression patterns of these genes. A small group of genes with an opposite expression pattern and high expression in the G1 phase were also observed. Nevertheless, we did not observe an obvious time-course expression pattern for the upregulated DEGs after IAA treatment (data not shown). Considering that ribosomal proteins are specific to the G1 phase and other organelle genes are dominantly expressed in the S/M/C phase, the greater accumulation of downregulated organelle genes and the reversed gene alteration for ribosomal subunit genes at 24 h post-treatment compared to 12 h of IAA treatment (Figure 3G and H) may be due to the increased proportion of parasites in the G1 phase as a result of G1 phase arrest following TgAP2XII-8 depletion.

For the 662 genes upregulated after 12 h of treatment, 31% (206/662) genes overlapped with the enteroepithelial developmental stages (EES)-specific genes (Figure 3I, Supplementary table 2), such as merozoite marker GRA82. The EES-specific genes (180/709) were also enriched in the upregulated genes after 24 h of IAA treatment (Figure 3J, Supplementary table 3). These results prompt a depletion of MORC, TgAP2XII-1 and TgAP2XI-2 [47], which suppress merozoite gene expression in the tachyzoite stage, and whose depletion leads to pre-sexual merogony development.

### 3.4 TgAP2XII-8 targets promoters of ribosomal proteins and beyond

To identify the direct binding profile of TgAP2XII-8 on the *T. gondii* genome, Cleavage Under Targets and Tagmentation (CUT&Tag) experiments were performed using endogenously 3×HA tagged TgAP2XII-8-mAID tachyzoites and RH strains. A total of 3378 peaks were identified in both biological replicates (*P* < 0.0001), of which 3180 peaks were assigned to the nearest genes and 95% of the peaks (3040) were located within 2 kb distance to the transcription start site (TSS) (Figure 4A and B). However, by overlapping these potential binding targets of TgAP2XII-8 with the RNA-seq DEGs, we obtained 335 and 443 genes that were potentially targeted by TgAP2XII-8 and differentially transcribed after 12 and 24 h of IAA-induced TgAP2XII-8 depletion, respectively (Figure 4C). Analogous gene numbers were found in the downregulated (47% for 12 h and 51% for 24 h DEGs) or upregulated parts of these targets (Figure 4D, Tables S1 and S2), which may suggest both activating and suppressing effects for TgAP2XII-8.

**Figure 4.**
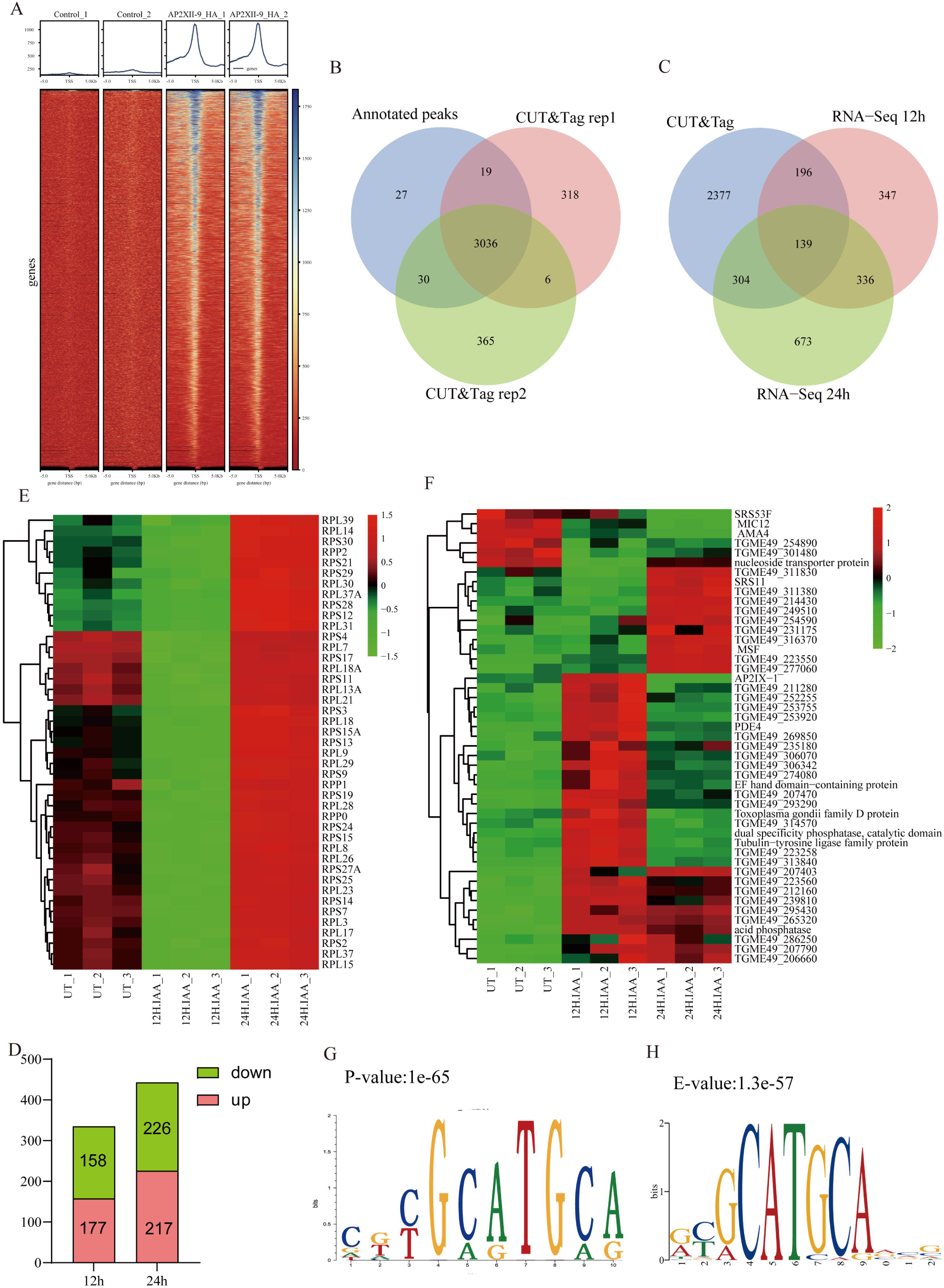
TgAP2XII-8 targets promoters of ribosomal proteins and beyond. (A) CUT&Tag analysis of TgAPXII-8 binding profiles around gene TSS. Upper panels: Average signal profiles centered on the TSS (±5 kb); Lower panels: Heatmaps representing the density of TgAPXII-8 binding peaks in the vicinity of the TSS. The color gradient on the right of each panel corresponds to the relative signal intensity. (B) Venn diagram showing the repeat experiments and annotation results. (C) Venn diagram showing the overlap of CUT&Tag hits and DEGs after TgAPXII-8 depletion at different times. (D) Statistics of CUT&Tag hits for downregulation and upregulation in the two RNA-SEQ datasets. Red sections represent upregulated target genes and green sections represent downregulated target genes. (E) Clustered transcriptional heatmap showing CUT&Tag hits of ribosomal proteins differentially expressed after TgAP2XII-8 depletion for 12 h and 24 h. (F) Clustered transcriptional heatmap showing CUT&Tag hits of enteroepithelial stage-specific genes differentially expressed after TgAP2XII-8 depletion for 12 h and 24 h. (G, H) TgAP2XII-8 DNA binding motif enrichment analysis using Homer (G) and MEME (H) algorithms. Statistical significance was marked.

A further analysis of the potential target genes of TgAP2XII-8 revealed that all 31 ribosomal proteins were downregulated after 12 h of IAA treatment, whereas 21 ribosomal proteins appeared to be upregulated after 24 h of treatment (Figure 4E, Supplementary figure 1). Since we hypothesized that the proportion of parasites at the G1 phase increased in the RNA-seq sample as a result of G1-phase arrest caused by TgAP2XII-8 depletion, we explain the upregulation of ribosomal proteins after 24 h of IAA treatment, which is a result of asymmetric sampling (which is unavoidable for the bulk RNA-Seq). As such, ribosomal proteins were downregulated immediately after TgAP2XII-8 degradation, which is also supported by the results of Luo *et al*. sampled only after 2 h of IAA treatment[48]. These results further suggest that TgAP2XII-8 acts as an activator for these ribosomal proteins.

Apart from these ribosomal proteins, TgAP2XII-8 may also control the expression of other important genes. There were 24 IMC and apical complex genes hit by CUT&Tag, of which 22 were downregulated even in 12 h of IAA treatment (45/46 for 24 h of treatment, Tables S1 and S2, Supplementary figure 1). The downregulation of these genes probably suffering from G1-phase arrest as explained above, however, these results may also suggest that TgAP2XII-8 has an activation role on these genes. On the other hand, TgAP2XII-8-targeted EES genes were upregulated by 83% (29/35) and 91% (33/36) at 12 and 24 h of IAA treatment, respectively (Figure 4F, Supplementary figure 1), revealing a suppressive effect of these genes.

The DNA-binding motif of TgAP2XII-8 was enriched by sequences under CUT&Tag peaks by using both MEME and HOMER, and both algorithms identified a similar motif ([T/C]GCATGCA) with high confidence (Figure 4G and H). This motif is highly consistent with the previously reported promoter element *<u>T</u>oxoplasma*<u>R</u>ibosomal <u>P</u>rotein (TRP)-2 ([T/C]GCATGC[G/A]), which is present in a majority of the upstream sequences of ribosomal proteins[49]. However, the other ribosomal protein promoter element TRP1 was not enriched in this study (data not shown). A further screening of the occurrence of the identified motif in upstream sequences (2 kb) of all *T. gondii* protein-coding genes by FIMO revealed that the motif was localized at an average distance of -1007 bp from the ATG codon of 5805 genes (data not shown). Over 3000 protein-coding genes were identified as binding targets of TgAP2XII-8, and more than 5800 genes had upstream sequences with binding potential, but only a small subset of genes had altered expression upon TgAP2XII-8 depletion (Tables S1 and S2). Together, these results suggest that other factors are required for the function of TgAP2XII-8 or are essential for the regulation of genes harboring this motif. In general, our results suggest that TgAP2XII-8 not only targets ribosomal proteins, but also controls the expression of many other proteins.

### 3.5 TgAP2XII-8 knockdown leads to division defects

The IMC is an important organelle in *T. gondii* that plays critical roles in parasite motility, invasion, egress, and replication[50, 51]. The IMC of *Toxoplasma* extends along the length of the parasite’s periphery and is composed of two layers: a series of flattened vesicles called alveoli and a rigid cytoskeletal network that supports intermediate filaments[52, 53]. As mentioned above, we found that several IMC-related genes may be targets of TgAP2XII-8. Generally, depletion of TgAP2XII-8 resulted in the downregulation of 33 IMC proteins after 12 h of IAA treatment, including IMC1, IMC3-7, IMC10, IMC12, IMC17, IMC26, IMC33, IMC37, IMC42, DHHC2, DHHC14, GAP40, GAP45, GAP50, GAPM2a, GAPM2b, GAPM3, AC3, AC4 and ERK7 (Supplementary table 2). After 24 h of IAA treatment, 87 IMC proteins were downregulated (Figure 5A). Notably, based on protein location, IMCs can be further divided into the apical cap (AC) at the top of the parasite, the centrosome portion, and the circular basal complex at the base of the parasite[54]. Among the downregulated transcripts, we unexpectedly found that deletion of TgAP2XII-8 induced significant downregulation of 18 basal complex genes after 24 h of IAA treatment, including BCC1, BCC2, BCC3, BCC4, BCC5, BCC6, BCC7, BCC9, BCC10, BCC11,

**Figure 5.**
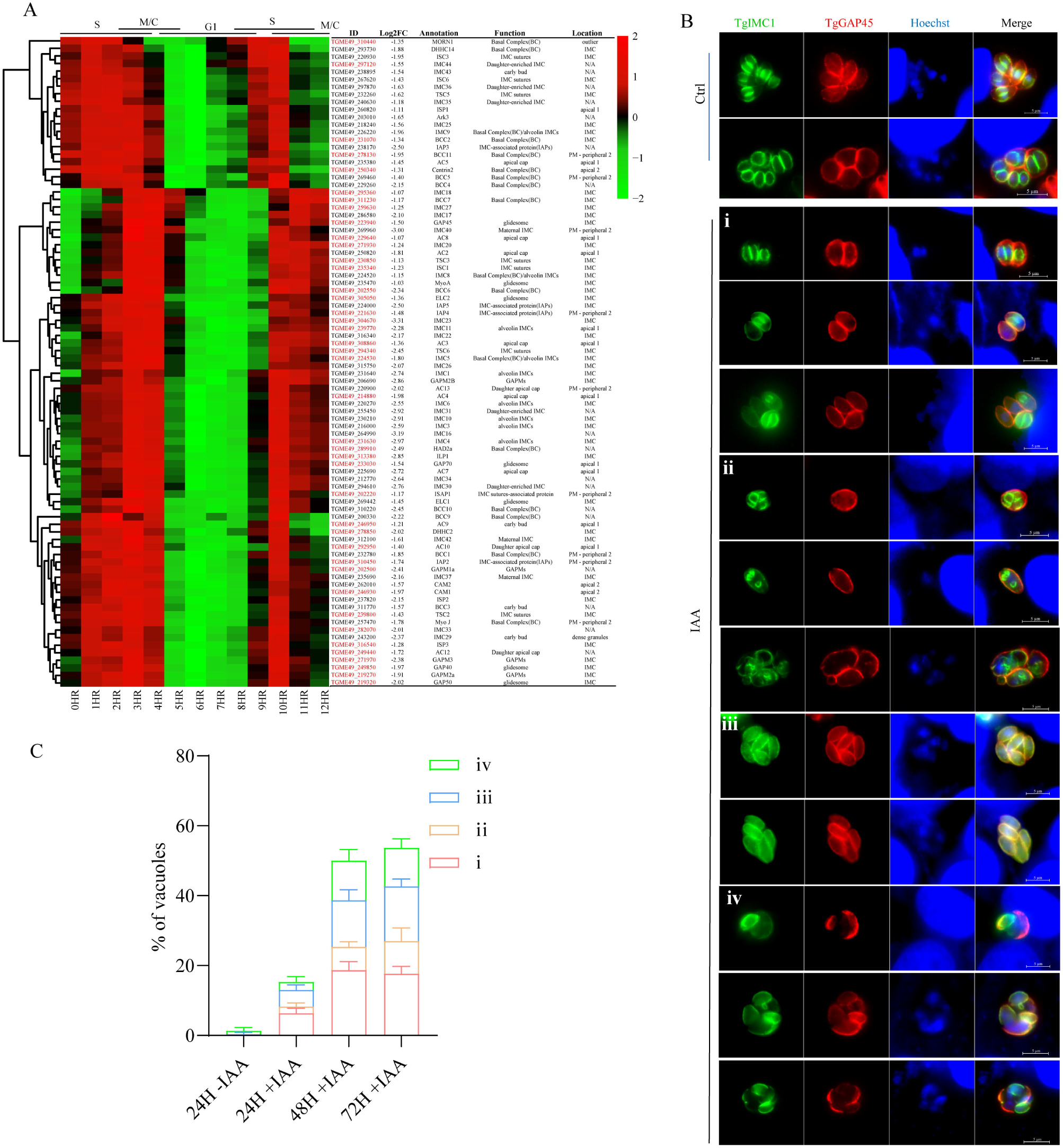
Disruption of TgAP2XII-8 leads to division defects. (A) Clustered heatmap of cell cycle expression of the differently expressed 87 IMC-related proteins, including Basal complex (BC) proteins, alveolin domain containing IMC proteins, gliding-associated membrane proteins (GAPM), glideosome-associated proteins, and IMC suture proteins. CUT&Tag hits are marked in red. (B) Immunofluorescent assay showing division defects after TgAP2XII-8 depletion. iKD TgAP2XII-8 parasites were treated with IAA or vehicle for 24 h, 48 h or 72 h. Fixed parasites were labeled against anti-TgIMC1 and anti-TgGAP45 antibodies. Scale bar = 5μm. i) non-synchronous division; ii) endopolygeny with >2 daughter buds per maternal parasite; iii) the presence of an odd number of tachyzoites in a PV; and iv) broken parasitophorous vacuoles. (C) Quantification of the above four types of abnormalities from three experiments (n□=□100 vacuoles). Asterisks indicate statistical significance. ** *P* < 0.01, *** *P* < 0.001.

Centrin2, DHHC14, HAD2a, MORN1, MyoJ, IMC5, IMC8, and IMC9. Among these downregulated genes, the promoters of IMC4, IMC5, IMC7, IMC12, IMC33, DHHC2, GAP40, GAP45, GAP50, GAPM2a, GAPM3, BCC2, BCC6, BCC7, and Centrin2 were bound by TgAP2XII-8, as shown in the CUT&Tag data. IMC7 and IMC12[55, 56] are recruited to the cytoskeleton outside of cytokinesis during the G1 cell cycle stage and are required for stress resistance of extracellular tachyzoites and maintenance of stability in mature parasites. Spatiotemporal localization kinetics analysis revealed that BCC6[57] and BCC7[58] are exclusively localized to the basal complex of mature parasites following the completion of cell division. The expression pattern of these four genes is consistent with that of TgAP2XII-8, suggesting that they may be directly activated by TgAP2XII-8 and that TgAP2XII-8 plays a role in daughter cell maturation.

However, as shown in the heatmap based on cell cycle data (Figure 5A), most of these genes are highly expressed during the S/M phase and reduce immediately after the C phase, whereas TgAP2XII-8 is expressed in the C/G1 phase as determined by IFA. Additionally, the CUT&Tag peaks and DNA-binding motifs of TgAP2XII-8 are widely distributed across the promoters of *T. gondii* genes, which are not only expressed during the C/G1 phase. Thus, we hypothesized that TgAP2XII-8 might act as a suppressor of these genes expressed in the S/M phase and initiate the cell cycle transition from the budding to the gap phase. However, these S/M-specific genes (e.g., IMCs) were found to be downregulated upon TgAP2XII-8 depletion, which is inconsistent with our hypothesis. This should be explained by the first downregulation of ribosomal proteins after TgAP2XII-8 depletion, which leads to G1-phase arrest in the parasites. Consequently, the read counts for genes that are highly transcribed in the S/M phase are relatively low due to the lower proportion of S/M-phase parasites in the bulk RNA-seq samples. On the other hand, the possibility of other interacting factors for TgAP2XII-8 cannot be excluded.

Given that several IMC proteins related to the formation of daughter parasites and IMC assembly are targeted by TgAP2XII-8, we further used TgIMC1 as a marker to verify whether the division of TgAP2XII-8-deficient parasites was affected. After treatment with IAA for 24 h, 48 h, and 72 h, iKD TgAP2XII-8 parasites exhibited a disorder of daughter division (Figure 5B). Four different types of unnatural divisions were mainly observed: i) non-synchronous division characterized by different budding states (budding, no budding, just budding, and complete daughter cell formation) of tachyzoites in the same PV; ii) endopolygeny with more than 2 daughter buds within a parasite (>2 daughter buds per maternal parasite); iii) the presence of an odd number of tachyzoites in a PV; and iv) broken parasitophorous vacuoles. The total proportion of abnormal divisions was 15.3% after 24 h of IAA treatment, whereas it increased significantly to 50% after 48 h of exposure (Figure 5C). However, the proportion of abnormally dividing parasites no longer increased after 72 h of IAA treatment (Figure 5C). In some cases, the odd number of tachyzoites observed in the PV was a consequence of non-synchronous divisions and the predominance of both types of unnatural divisions (62%-70% among unnatural divisions) after TgAP2XII-8 depletion. Our findings indicate a significant deficiency in the cytoskeleton assembly and daughter cell division after AP2XII-8 knockdown.

## 4 Discussion

Toxoplasmosis poses significant public health threats, with about one-third of the global population estimated to be infected. Acute infection is dependent on the rapid replication of *T. gondii* tachyzoites inside host cells. *T. gondii* tachyzoites undergo successive coordinated binary divisions within the parasitophorous vacuole, during which thousands of genes are precisely expressed in different cell-cycle phases. Previous studies have shown that 24 ApiAP2 transcription factors are cyclically expressed during tachyzoite division. However, the exact function and targets of several ApiAP2s remain unknown. Here, we show that the C/G1 phase-expressed nucleotide transcription factor TgAP2XII-8 is crucial for parasite intracellular replication. TgAP2XII-8 targets the ribosomal subunit genes and several IMC/apical genes, and depletion of TgAP2XII-8 results in significant G1 phase arrest and division disorders. Our results provide important insights into the G1 phase gene transcriptional regulation in *T. gondii*.

The pathogenesis of acute *T. gondii* infection is largely related to the rapid asexual amplification of tachyzoites inside host cells. The asexual reproductive cell cycle of *T. gondii* tachyzoites consists of five major phases, termed G1, S, G2, M and C (Cytokinesis or budding), with the recent identification of the conventional G2 phase preceding the M phase[5]. G1 phase is usually conserved between species and is the longest period responsible for preparing materials for the next replication cell cycle[10]. RNA processing and protein translation-related proteins (e.g., ribosomal proteins) are synthesized mainly in the early G1 phase (G1a), whereas DNA replication and repair-related proteins are produced mainly in the late G1 phase (G1b)[8]. In this study, we found that TgAP2XII-8 was expressed during G1 phase, and the degradation of TgAP2XII-8 resulted in the downregulation of multiple ribosomal proteins but limited alterations in DNA replication/repair-related proteins. These results may indicate that TgAP2XII-8 acts as an activator of gene transcription in G1a but not in G1b. Downregulation of ribosomal proteins after TgAP2XII-8 degradation leads to breakdown of protein synthesis in the G1a phase, which disturbs the parasite’s entry into the next cell cycle and results in G1-phase arrest. Additionally, TgAP2XII-8 was found to be expressed in C phase when daughter cell assembly is nearing completion. Our results demonstrate that TgAP2XII-8 targets not only ribosomal genes, but also IMCs and apical/basal complex genes, and that depletion of TgAP2XII-8 leads to a disorder of parasite budding and daughter cell formation, suggesting that TgAP2XII-8 also plays a very important role in the cytokinesis phase of *T. gondii* tachyzoites.

In this study, we found that several IMC and basal complex genes were significantly downregulated after TgAP2XII-8 depletion, and their promoter sequences were proved to be bound by TgAP2XII-8. Usually, TgAP2XII-8 can accordingly be considered as an activator for these genes. However, TgAP2XII-8 was demonstrated to be expressed during the C to G1 phases, while most of these IMC and basal complex genes exhibit expression peaks in the S/M phase and a rapid decline during the C phase, with the exception of IMC7, IMC12, BCC6 and BCC7, which was expressed in parallel with the expression pattern of TgAP2XII-8. The expression pattern of these IMCs and TgAP2XII-8 is reminiscent of the fact that TgAP2XII-8 acts as a suppressor for these IMCs and as a terminator of the division process by suppressing such genes that are specifically expressed in the S/M phase. The downregulation of these IMCs is attributable to the bulk sampling of RNA-seq and G1-phase arrest following TgAP2XII-8 depletion. Ribosomal proteins also showed this interference, as they were downregulated 12 h after TgAP2XII-8 depletion but upregulated 24 h later. Thus, our data support the idea that TgAP2XII-8 acts as key switch in the transition from the S/M phase to the C phase by acting as a suppressor of highly expressed genes during the S/M phase (e.g., IMC genes) and as an activator of G1 phase-specific genes (ribosomal proteins).

By CUT&Tag enrichment, we found that TgAP2XII-8 binds to a specific DNA motif ([T/C]GCATGCA) on the *T. gondii* genome. This motif is identical to the TRP2 *cis*-element localized in the 5’-untranslated region of most ribosomal protein genes in *T. gondii*[49], which suggests a regulatory role for TgAP2XII-8 on ribosomal protein genes. The motif was also found to be extensively represented in hundreds of promoters flanking the G1-subtranscriptome genes, while underrepresented in promoters flanking genes of the S/M-subtranscriptome, as predicted by FIRE (Finding Informative Regulatory Elements) analysis using the 2kb 5’UTR of *T. gondii* genes [8]. The motif is not restricted to the promoters of ribosomal protein genes expressed in the G1 phase, but is widely dispersed among the >5000 flank sequences of the *T. gondii* protein-coding genes. Considering that the transcription of only some of the genes is altered after TgAP2XII-8 depletion, we hypothesized that there are other factors that are essential for the regulation of genes harboring this motif. This hypothesis is supported by the similar DNA-binding motif of TgAP2XI-3 and TgBDP1 (bromodomain protein) [8, 59] TgAP2XI-3 is expressed at its peak in the G1 phase similar to TgAP2XII-8, and may co-opts with each other. The potential direct interaction or binding of TgAP2XII-8 and TgAP2XI-3 as a complex should be further validated. The lysine acetylation reader TgBDP1 binds to the promoters of large number of genes, and show both activation and suppression role for the expression of these genes[59]. However, the interactome of TgBDP1 did not support a direct interacting between TgBDP1 and TgAP2XII-8[59], suggesting an independent regulation of *T. gondii* genes with [T/C]GCATGCA motif by TgBDP1 and TgAP2XII-8.

The motif bound by TgAP2XII-8 is very close to a palindromic octamer TGCATGCA, which is not only abundant and conserved in the genome of *T. gondii*, but also in the intergenic regions of several other apicomplexans, including *P. falciparum*, *Cryptosporidium parvum*, and *Eimeria tenella*[60]. This motif binds to the *C. parvum* transcription factor cgd2_3490 and has also been characterized as the binding motif for PfAP2-EXP and its rodent orthologue PbAP2-SP[61]. Binding sites for PfAP2-EXP have been predicted to be located upstream of members of the *var* gene family and other ApiAP2 genes[61, 62], and PfAP2-EXP is essential for the *P. falciparum* asexual blood stage[63]. PbAP2-Sp is dispensable for asexual parasite growth and is a master regulator of sporogony, and the motif TGCATGCA, to which PbAP2-Sp binds, was validated as a *cis*-acting element specific for the sporozoite stage in *P. berghei*[64]. Additionally, the palindromic octamer TGCATGCA motif matches almost perfectly with the nucleosome-around sequences in *P. falciparum*, especially at later stages of parasite development[65]. Considering the overrepresented distribution and palindromic nature, this motif is expected to play a role in chromatin packaging or dynamics[60] and suggests the existence of an evolutionary conserved regulatory pattern in apicomplexan parasites. More importantly, upon dissecting the X-ray crystal structure of the PfAP2-EXP AP2 domain[66], a set of chemical compounds targeting its AP2 DNA-binding domain were identified, which are highly promising for new classes of antimalarial therapeutics[67]. These advances in malaria shed light on drug discovery and control of toxoplasmosis.

In conclusion, this study characterizes a C/G1 phase that expresses the ApiAP2 transcription factor TgAP2XII-8, which acts as an activator for ribosomal protein genes and possibly a suppressor for genes that are highly expressed in the S/M phase. We demonstrate that the depletion of TgAP2XII-8 results in G1 phase arrest and division disorder. Our study provides crucial data for the understanding the transcriptional regulation of the cell cycle in *T. gondii* parasites.

## AUTHORS CONTRIBUTIONS

Conceptualization, Formal analysis, Funding acquisition, Project administration, Resources, Supervision, review and editing the manuscript, D.H. and X.S.; Data curation, Formal analysis, Investigation, Methodology, drafted the manuscript, Y.S.; Investigation, Methodology, Validation, Q.Y. and Y.H.

## ACKNOWLEDGMENTS

We are grateful to Prof. Qun Liu, Prof. Jing Liu and Prof. Shaojun Long (China Agricultural University, China) for providing parasite strains and antibodies. Prof. Dongying Wang, Yucong Jiang, Mao Huang, Xinru Cao and Yazhen Ma in our institution are also acknowledged for their technical help.

## DATA AVAILABILITY STATEMENT

The data that support the findings of this study are available in the methods and / or supplementary material of this article. RNA-Seq data generated in this study have been deposited to the NCBI Sequence Archive (SRA) under the accession number PRJNA1148922. Raw sequencing data and processed data for CUT&Tag experiments are available in the NCBI GEO database under the accession number GSE275112.

## CONSENT TO PUBLISH

All authors have approved the content of this manuscript and provided consent for publication.

## SUPPORTING INFORMATION

**Supplementary table 1:** Primers used in this study.

**Supplementary table 2:** Integrated CUT&Tag hits and transcriptomic data for the iKD TgAP2XII-8 strain after 12 h of IAA treatment.

**Supplementary table 3:** Integrated CUT&Tag hits and transcriptomic data for the iKD TgAP2XII-8 strain after 24 h of IAA treatment.

**Supplementary figure 1:** IGV snapshots of the genome regions of represented genes based on RNA-seq and CUT&Tag data.

