## Supplementary figures and images for "*Toxoplasma gondii* transcription factor AP2XII-8: a key regulator of G1 phase progression and parasite division"

### supplementary Figure 1.pdf

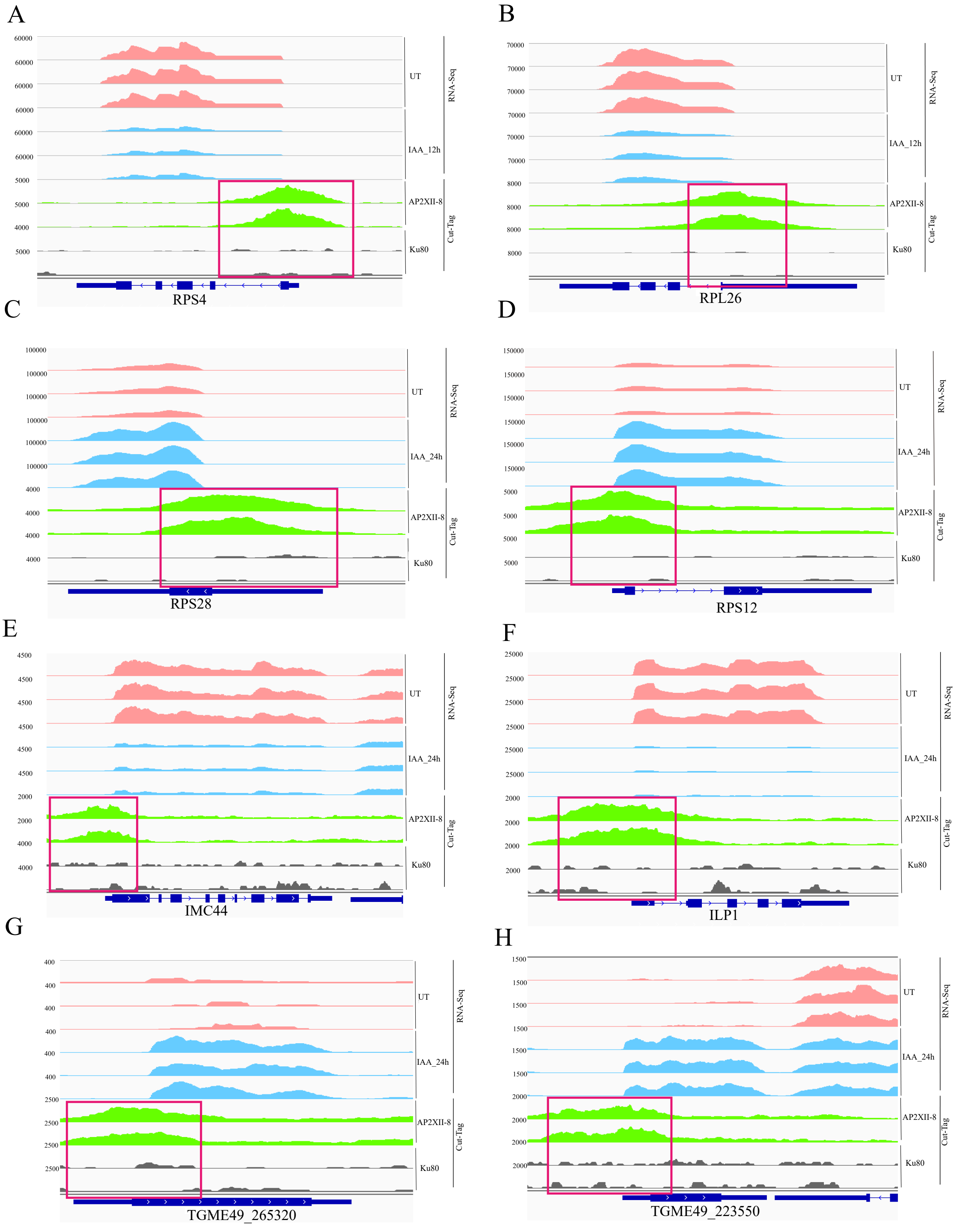
